# Structure of the human pro-myostatin precursor and determinants of growth factor latency

**DOI:** 10.1101/153403

**Authors:** Thomas Cotton, Gerhard Fischer, Xuelu Wang, Jason McCoy, Magdalena Czepnik, Thomas B. Thompson, Marko Hyvönen

## Abstract

Myostatin, a key regulator of muscle mass in vertebrates, is biosynthesised as a latent precursor in muscle and is activated by sequential proteolysis of the pro-domain. To investigate the molecular mechanism by which pro-myostatin remains latent, we have solved the structure of unprocessed promyostatin and analysed the properties of the protein in its different forms. Crystal structures and SAXS analyses show that pro-myostatin adopts an open, V-shaped structure with a domain-swapped arrangement. The pro-mature complex, after cleavage of the furin site, has significantly reduced activity compared with mature growth factor and persists as a stable complex that is resistant to the natural antagonist follistatin. The latency appears to be conferred by a number of distinct features that collectively stabilise the interaction of the pro-domains with the mature growth factor, enabling a regulated step-wise activation process, distinct from the prototypical pro-TGF-β1. These results provide a basis for understanding the effect of missense mutations in pro-myostatin and pave the way for the design of novel myostatin inhibitors.

## INTRODUCTION

The pathological outcomes which arise as a result of aberrant cellular signalling, including cancer, highlight the importance of spatial and temporal signal control in biology. One of the ways that signalling protein activity can be controlled is by the expression of these molecules as inactive, or latent forms, with activation occurring only where and when a timely response is required. Controlled proteolysis is a common mechanism of activation for the pro-forms of bioactive molecules and is well characterised in many biological systems, from the proteases of the digestive system, to secreted growth factors. Controlled post-translational activation allows the proteins to be expressed and stored in a precursor form and then rapidly activated in response to external stimuli.

Myostatin, (also called growth and differentiation factor 8; GDF8), of the transforming growth factor β (TGF-β) superfamily of signalling proteins is a negative regulator of skeletal muscle growth. Dysfunctional myostatin signalling liberates muscle growth and yields the characteristic hyper-muscular phenotypes seen in myostatin-null animals^1,2^. Unsurprisingly, manipulation of myostatin signalling has become an attractive prospect for increasing functional muscle mass in the context of muscular atrophic disorders including muscular dystrophy, sarcopenia and cancer-associated cachexia^3^.

Myostatin itself is a relatively well characterised member of the TGF-β superfamily, and like other members, is synthesised as an inactive precursor (pro-myostatin), with N-terminal signal peptide and pro-domain, and C-terminal growth factor (GF) domain. The precursor forms a covalently linked dimer through a conserved disulfide in the GF domain^1,4,5^. Cleavage of pro-domains by furin-like pro-protein convertases, either during secretion or extracellularly, yields a non-covalent complex of the dimeric mature GF with its associated pro-domains (pro-myostatin complex)^6,7^. The non-covalent association of pro-domains is typically thought to retain myostatin in a latent state by occluding receptor epitopes and rendering it unable to engage its receptors^5,6^. In contrast to pro-TGF-β1 which undergoes integrin-driven mechanical activation, a secondary proteolytic cleavage within the pro-domain by BMP1/Tolloid (TLD) family metalloproteases liberates the full signalling capacity of mature myostatin^6,8^. The liberated, mature myostatin will form a heterotetrameric complex with two activin responsive type II receptors (ActIIRA or ActIIRB) and two of either activin type I (ALK4) or TGF-β type I (ALK5) receptors to initiate signalling^4,9^. Assembly of a competent receptor complex results in SMAD 2/3 phosphorylation by the type I receptors and translocation of SMADs to the nucleus for modulation of gene expression^10^.

At present, three structures of pro-TGF-β superfamily members are available; pro-TGF-β1, pro-activin A and pro-BMP9, all of which display unique arrangements of pro and mature GF domains^8,11,12^. As mentioned above, in some cases the pro-domain confers latency to the pro-form, as is known to be the case for pro-TGF-β1 and pro-myostatin. Conversely, pro-activin A and pro-BMP9 complexes show equivalent signalling activity to their free mature GFs, suggesting a weak, non-inhibitory association of pro-domains.^11,12^ TGF-β1, which forms a latent complex with its furin-cleaved pro-domains, utilises an inter-molecular disulfide bond to cross-link pro-domains and enclose the dimeric GF in an inhibitory stranglehold, requiring mechanical or proteolytic activation^8^. Pro-myostatin lacks the cysteines needed for this this latency conferring covalent linkage, and as such the structural basis for its latency remains unclear.

Myostatin is known to be secreted both as an unprocessed precursor and a furin cleaved complex, with the former thought to constitute the major pool of myostatin in the extracellular space of skeletal muscle^13^. Within the extracellular environment of muscle, latent pro-myostatin is localised to the extracellular matrix (ECM), through pro-domain mediated interactions with heparan sulfate proteoglycans and latent TGF-β binding proteins (LTBPs)^13,14^.Soluble antagonists of the mature GF, including follistatin, FSTL3, GASP1, GASP2 and decorin, contribute an additional layer of control, within an already complex regulatory environment^15–17^.

Targeted inhibition of myostatin signalling to enhance muscle growth continues to present a considerable clinical challenge. A number of myostatin binding antibodies, designed to suppress myostatin signalling in the context of muscular atrophic disorders, have failed to meet primary clinical endpoints in phase II trials (bimagrumab, Novartis; PINTA 745, Atara)^18,19^. Similarly, an ActRIIB receptor-Fc fusion (ACE-031, Acceleron) was withdrawn from phase II trials due to safety concerns^3^. To date, no myostatin inhibitors are approved for clinical use. It seems probable that attempts to block mature myostatin signalling are hampered by the cross-reactivity of soluble antagonists with structurally related TGF-β superfamily growth factors. This is particularly likely for soluble receptor-Fc fusions, or ligand-traps, which exhibit a natural promiscuity towards different TGF-β superfamily ligands.

The difficulties associated with targeting the mature, active growth factor, make the more structurally diverse pro-forms of these proteins potentially more meaningful targets for intervention. Aside from lower conservation in sequence and structure, the pro-forms show increased abundance and longevity over the mature growth factors which are difficult to target due to the short temporal and spatial window within which they exert their paracrine signal. Stabilisation of a latent conformation of the pro-myostatin complex and/or inhibition of proteolytic processing of the precursor could offer alternate routes to selective neutralisation of myostatin signalling. Understanding the mechanism by which the pro-domains render the growth factor inactive will be essential for these efforts.

Here we present crystal structures of unprocessed human pro-myostatin, the major extracellular storage form in skeletal muscle tissue. This structure reveals a unique arrangement of GF and pro-domains to confer latency within the TGF-β superfamily. An unexpected ‘open-armed’ conformation, with no direct interaction between the arm/shoulder-domains of the domain-swapped dimer, makes pro-myostatin structurally distinct from latent pro-TGF-β1. This structure allows us to understand the determinants of latency and reveals features that enable controlled activation of myostatin. It provides us also with a rational basis for the development of the next generation of myostatin inhibitors.

## RESULTS

### Production and characterisation of human pro-myostatin

Unprocessed human pro-myostatin was expressed as inclusion bodies in *E. coli* and subsequently solubilised, refolded and purified. As expected, the protein migrated on non-reduced SDS-PAGE as a disulfide-linked dimer and analysis by size exclusion chromatography and multi-angle light scattering (SEC-MALS) confirms the dimeric state under native conditions, with a molecular weight of 84.5 ± 0.005 kDa (cf. calculated from sequence 85.4 kDa) (Figure 1A).

**Figure 1.**
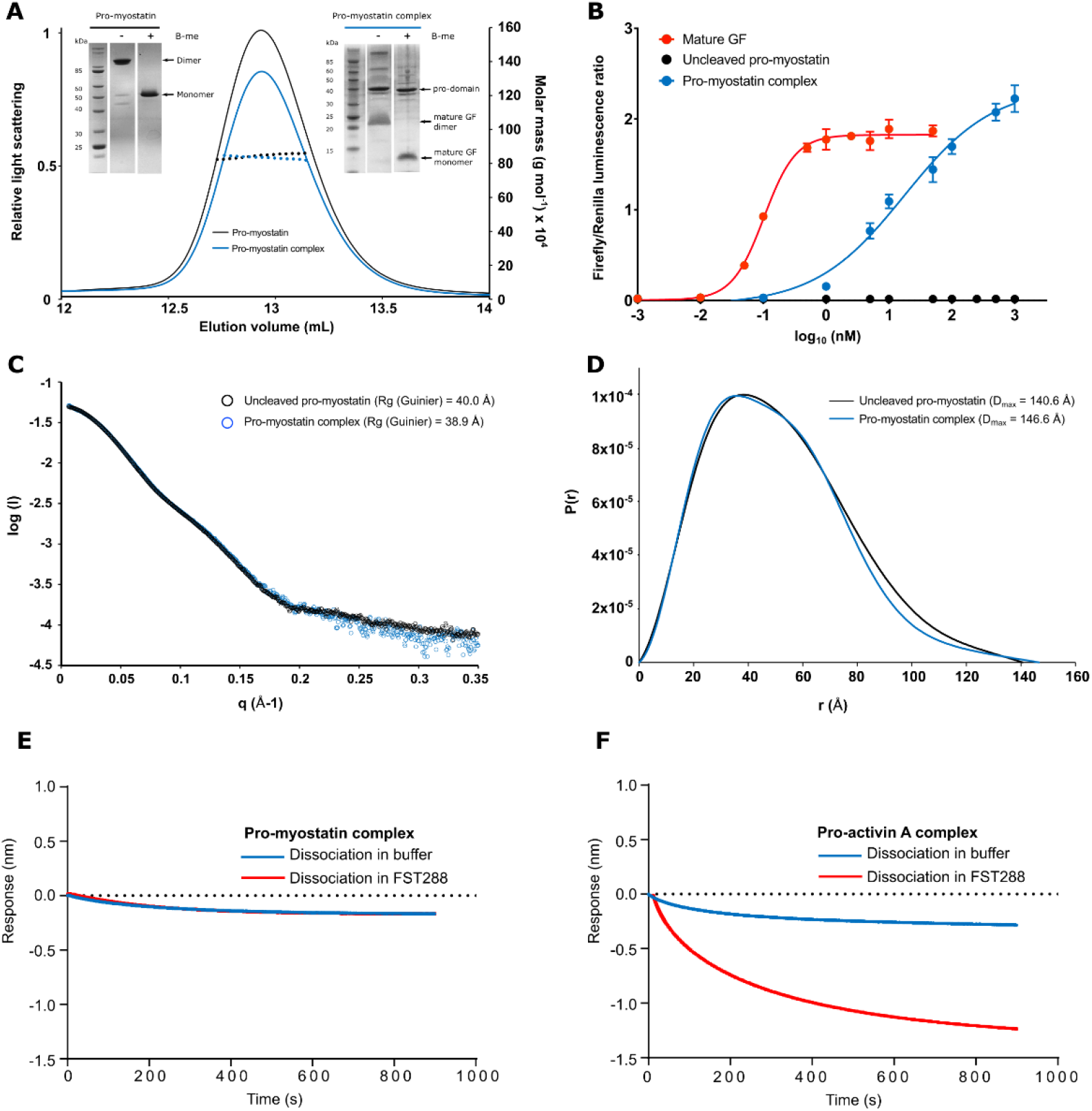
Characterisation of pro-myostatin in solution A. SEC-MALS analysis of pro-myostatin, and HRV-3C cleaved complex (loaded samples shown on inset gels). The cleaved complex elutes at same volume as the uncleaved precursor, indicative of stable complex formation between mature GF dimer and two pro-domains. B. Myostatin signalling response in HEK293T luciferase reporter assay. Purified mature GF domain is more than 100 times as potent as the ‘latent’ complex, while uncleaved pro-myostatin shows no signalling activity. C. Small-angle X-ray scattering intensity (I) vs q for pro-myostatin and HRV-3C cleaved complex. Scattering curves overlay well, with little change in the estimated radius of gyration (R_g_) following cleavage of the pro-domain. D. Inter-atomic pair distribution functions P(r) calculated by DATGNOM (q_max_ ≈ 0.2) for pro-myostatin and HRV-3C cleaved pro-myostatin complex. P(r) functions approach zero smoothly at D_max_=140.6 Å and 146.6 Å for uncleaved and cleaved pro-myostatin respectively. E. Dissociation of mature myostatin GF from cleaved pro-myostatin complex using biolayer interferometry. The complex is immobilised on sensor tip through N-terminal His-tag using an anti-penta-His antibody and dissociation of the mature domain monitored for 900 seconds in absence (blue line) and presence (red line) of 500 nM FST-288. F. Same experiment shown in E. but monitoring dissociation of pro-activin A complex.

For functional analysis, an engineered variant of pro-myostatin was generated in which the native furin cleavage site was replaced by a HRV-3C protease site, to allow us to study the effects of proteolytic processing of the precursor *in vitro*. The HRV-3C cleaved pro-myostatin was shown, by SEC-MALS, to form a stable non-covalent complex with a molecular mass of 83.4 ± 0.008 kDa, consistent with the expected mass for a complex of a mature GF dimer with two associated pro-domains (Figure 1A). Pro-domain cleavage appears to proceed via a semi-cleaved intermediate, a small proportion of which persists in the final preparation, even after incubation with a molar excess of protease. The mature GF dimer could be purified from the complex by reverse phase chromatography, and was shown to activate SMAD2/3 signalling in HEK293T cells with high potency (EC_50_: 0.1 nM, 95% C.I [0.09, 0.12]) (Figure 1B). While uncleaved pro-myostatin was entirely inactive at the highest concentrations of protein tested, the HRV-3C cleaved complex shows low-level signalling activity, more than 100-fold less potent than the purified mature GF (EC_50_: 17 nM, 95% C.I [11, 32]) (Figure 1B). Bioactivity of the pro-myostatin complex, traditionally thought to circulate as an entirely latent complex in serum, has been observed previously by Szlama *et al*, who interpret the unexpected activity as the result of partial dissociation of the pro-domains under assay conditions^20^. This is in clear contrast to both pro-TGF-β1 and pro-activin A complexes. Latent pro-TGF-β1 shows no activity under similar assay conditions, whereas the pro-activin A pro-domain exerts only a marginal inhibitory effect at the picomolar concentrations where the mature growth factor has been shown to be active.^8,12^

To evaluate whether cleavage at the furin-site causes significant conformational change of the protein, we analysed both uncleaved and cleaved pro-myostatins using small angle X-ray scattering (SAXS). The scattering profiles were very similar in both cases and the estimated radii of gyration (R_g-,uncleaved_=40.0 Å, R_g,cleaved_=38.9 Å) and maximum particle dimensions (D_max,uncleaved_=140.6 Å, D_max,cleaved_=146.6 Å) of the proteins are very similar (Figures 1C, 1D, S1). This suggests that no drastic re-organisation of the protein is triggered by the proteolysis of the furin site, consistent with the significant latency of this complex in bioassays.

To study the stability of the cleaved pro-myostatin complex further, we used biolayer interferometry (BLI) to monitor the dissociation of the mature GF from the pro-domain, which was immobilised on biosensor tips through N-terminal His-tag. We observe very slow dissociation of the GF, consistent with the low level of activity seen in cellular assays (Figure 1E). Interestingly, this dissociation cannot be facilitated by natural myostatin inhibitor follistatin (FST-288, the 288 amino acid isoform). This is in stark contrast to the pro-activin A complex, which readily dissociates in the presence of FST-288 (Figure 1F). On the other hand, same experiment using uncleaved pro-myostatin shows no significant difference in the presence or absence of FST-288, while uncleaved pro-activin A actually shows an increase in response when exposed to FST-288, suggesting an interaction with growth factor part of activin A, even before proteolytic cleavage releases the mature domain from its pro-domain (Figure S2). These results confirm that the pro-myostatin complex is highly stable, more so than pro-activin A. These data suggests also that regulation of myostatin by follistatin can only take place after the pro-domain has dissociated from the mature GF.

### Structure determination

To elucidate the molecular determinants of pro-myostatin latency, we crystallised the uncleaved precursor form of human pro-myostatin, with its native furin site intact. This protein crystallised readily in a number of conditions, yielding cubic crystals. These were used to determine the structure at a resolution of 4.2 Å, using experimentally determined phases from selenomethione-labelled protein (**Figure S3**). Merging data from multiple crystals (**Table S1**) significantly improved the quality of the electron density for this structure, so that we could trace most of the backbone and observe large side chains. However, we were unable to unambiguously assign the sequence of the pro-domain using this low resolution data.

Using a surface entropy reduction approach^21^, we identified a mutant form of pro-myostatin that crystallised in a new crystal form and diffracted to higher resolution. One symmetrical half of the low-resolution dimer was used as the search model for molecular replacement of a higher resolution dataset, and the structure was solved again, this time at 2.6 Å (Figure 2A). In both crystal forms, the asymmetric unit contains a single dimeric molecule. We were able to build 92% and 81% of residues of the two protomers into the electron density of the 2.6 Å structure.The remaining regions are disordered, with 27 and 62 residues missing from chains A and B respectively. The quality of electron density differs markedly between different parts of the two chains (**Figure S4**), and as such, our interpretations are based on the analysis of both chains of the higher resolution structure.

**Figure 2.**
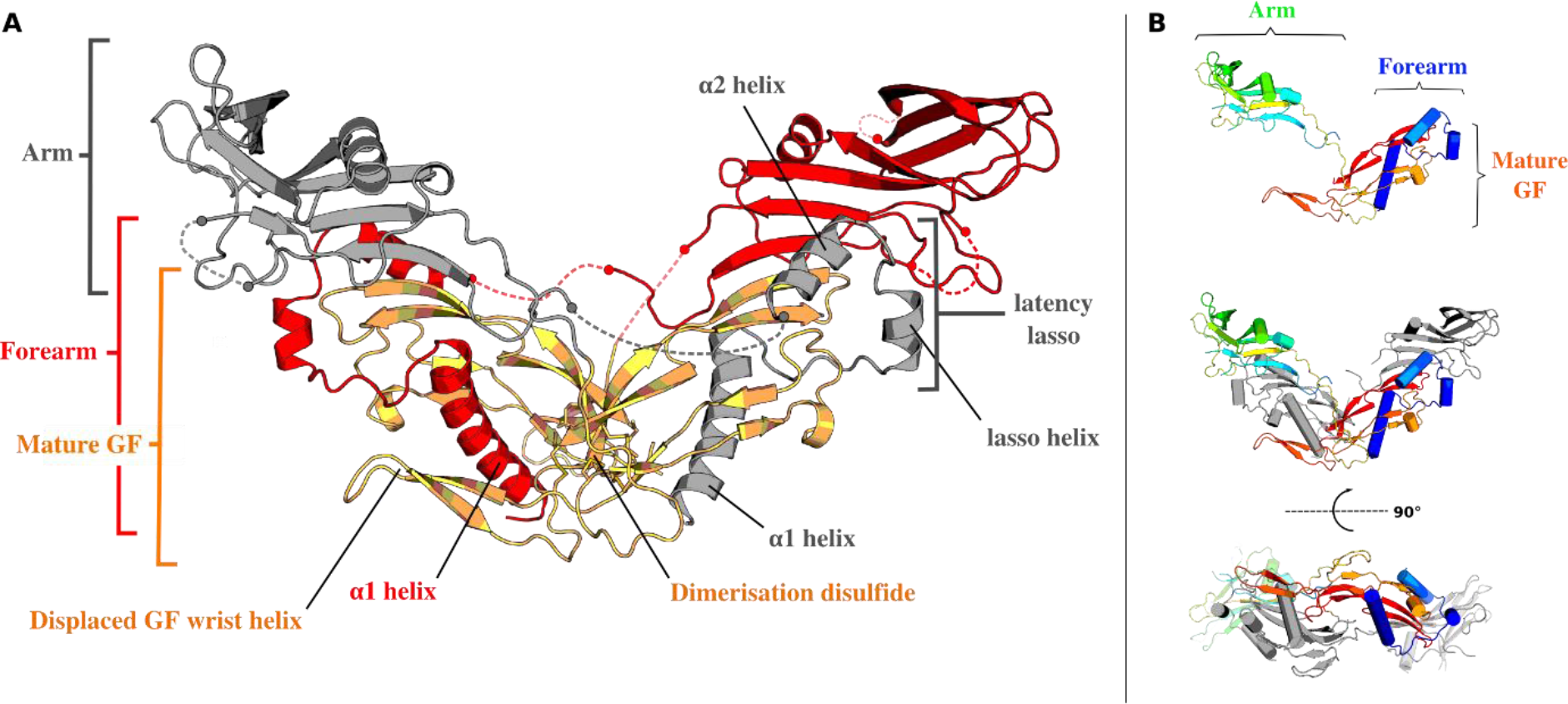
Structure of unprocessed human pro-myostatin. A. 2.6 Å structure of unprocessed pro-myostatin dimer showing mature GF dimer (orange) with bound pro-domains (red and grey). Unmodelled loop regions are shown as dashed lines. B. Pro-myostatin coloured by rainbow from N-terminus (blue) to C-terminus (red), for a single protomer (top) and dimer (middle and bottom, second protomer coloured grey).

### The structure of human pro-myostatin

Like related pro-TGF-β superfamily members, pro-myostatin is a disulfide-linked homodimer, each chain of which contains an N-terminal pro-domain and a C-terminal mature GF domain (Figure 2A). The GF domains consist of four antiparallel β-strands or ‘fingers’ and a cystine-knot motif, characteristic of TGF-β superfamily members. Two identical GF protomers associate through their concave ‘palms’, and are linked covalently through a disulfide bond between equivalent Cys339 residues in the GF ‘wrist’ region (Figure 2A). The pro-domain retains the familiar structural elements of other pro-TGF-β superfamily members, including N-terminal ‘forearm’ helices which grasp the mature GF, and a globular ‘arm/shoulder’ domain, which sits atop the mature GF protomers (Figure 2A).

Given the latency of the pro-myostatin complex, it was expected that the pro-domains would adopt a closed conformation like that of pro-TGF-β1, albeit without the cross-linking disulfide.^8^ Instead, pro-myostatin adopts a V-shaped, “open arm” conformation with no interactions between the arm domains, similar to that observed for the two non-latent complexes of pro-BMP9 and pro-activin A (Figure 3D).^11,12^

The individual chains of both our low and high resolution structures overlay well (Cα RMSD: 0.68 Å, 227 atoms, using non-covalently associated pro-and mature domains as a single entity). However, there is a considerable shift in the inter-protomer angle between the two structures, measured from the dimerization disulfide to the tips (Gln358) of the mature domain fingers, with the low resolution model adopting a more closed conformation (89.2° vs. 108.5°; Figure 3A). This suggests the pro-form has significant conformational flexibility about the dimer interface. To explore this, we used SAXS data to calculate a molecular envelope for uncleaved pro-myostatin. The resulting envelope shows an extended structure, consistent with what we see in the crystal structures, but is less well defined and multiple inter-protomer conformations, rotating about the dimerisation disulfide, could be accommodated within the envelope (Figure 3B).

**Figure 3.**
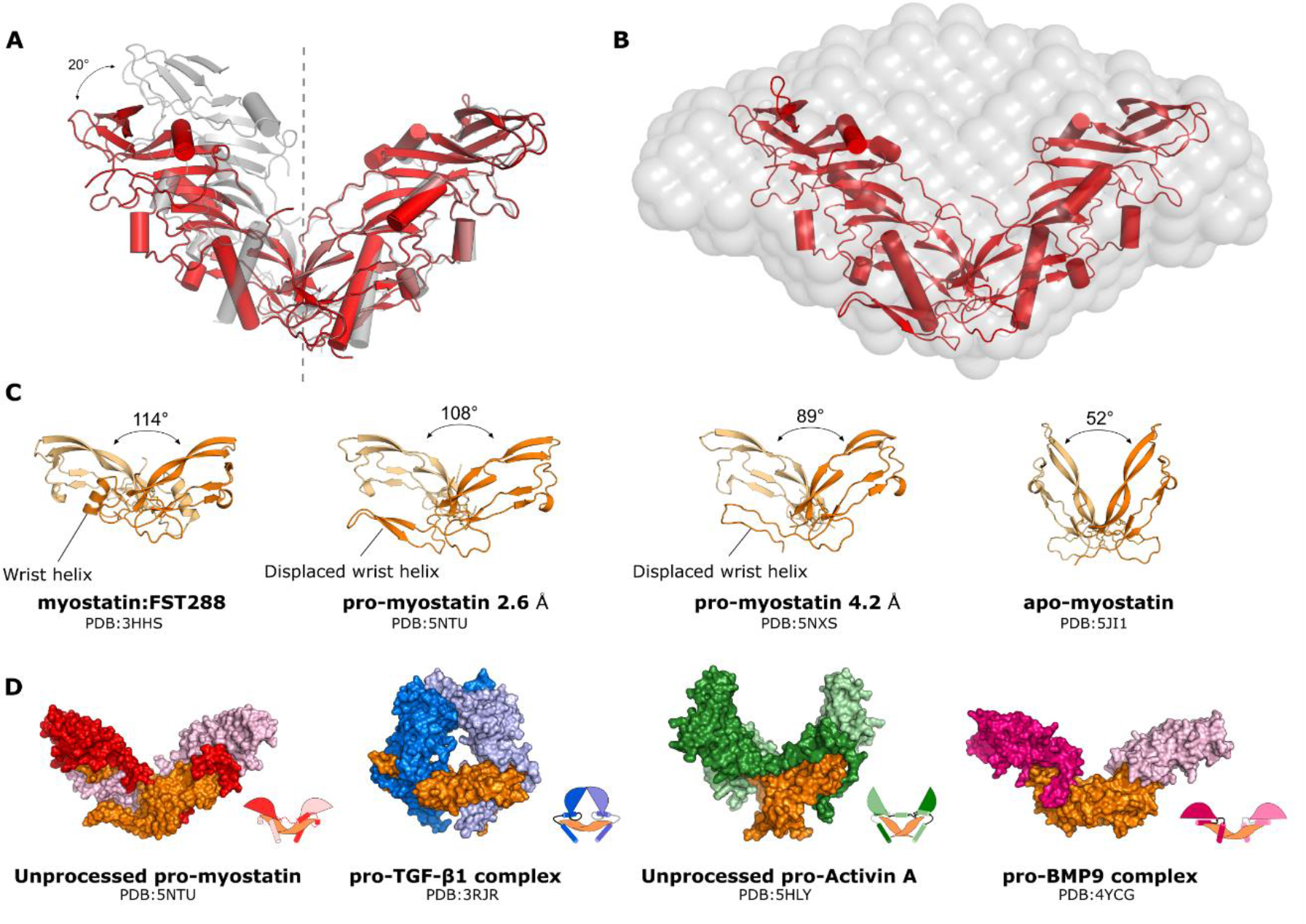
Conformational flexibility of pro and mature myostatin. A. High (red) and low (grey) resolution pro-myostatin structures aligned by a single mature GF protomer, showing shift in inter-protomer angle. B. *Ab initio* SAXS envelope (DAMFILT) of unprocessed pro-myostatin, with docked pro-myostatin structure (PDB: 5NTU). C. Mature myostatin GF dimers from structures solved to date, showing inter-protomer plasticity (individual protomers coloured orange and pale orange). D. Comparison of pro-myostatin with the architectures of known pro-forms of TGF-β superfamily growth factors.

The individual mature GF protomers also overlay well with the structure of myostatin bound to follistatin 288 (PDB: 3HH2, Cα RMSD: 0.63Å, 65 atoms), but exhibit a shift in inter-protomer angle (Figure 3C)^15^. This observation is consistent with that of Walker *et al*, who recently showed that the mature myostatin GF dimer crystallises with radically different inter-protomer angles in apo and FST-288 bound states (Figure 3C)^22^. Conformational plasticity is similarly well documented for activin A, which has inter-protomer angles ranging from 50° in complex with type II receptor ecto-domain (PDB: 1NYS), to 108° when bound to FST-315 (PDB: 2P6A)^12^.

In both of our structures, the GF domain ‘wrist’ helix and pre-helix region, which forms a significant interface with the opposing protomer, and establishes the presumed binding site for the type I receptor, is displaced in the presence of the pro-domain. Instead, this sequence forms a β–hairpin visible within the crystal contact of one chain and binds on the exposed face of the pro-domain α1 helix (Figure 2A). The GF wrist helix and pre-helix region are thought to constitute an important component of the putative type I receptor epitope and while there is currently no structure of a myostatin-receptor complex, the ALK5 binding mode can be inferred from the ALK5:TGF-β3 structure (2PJY)^23^. Displacement of the wrist helix by the pro-domain would render myostatin unable to engage the type I receptor while bound to its pro-domain. In contrast to this, the N-terminal domain (ND) of FST-288 occupies the type I receptor site without displacing the wrist helix, and this site is known to accommodate a number of different ligands by utilising this ‘non-invasive’ binding mode^15^. In pro-TGF-β1 the wrist helix is also displaced, and in pro-activin A density for the helix is missing altogether, suggesting similar displacement from the core of the GF domain^8,12^. In the pro-BMP9 structure, the wrist helix remains in place, with the α5 helix from the prodomain occupying a similar position to the helix of the FST-288 ND domain^11^.

### Pro-myostatin forms a domain-swapped, open armed dimer

Similar to previously determined structures of TGF-β family pro-domains, the myostatin pro-domain consists of an N-terminal α1 helix/loop/α2 helix ‘forearm’ motif, and a C-terminal globular ‘arm’ domain (Figure 2A)^8^. The pro-myostatin forearm is structurally similar to that of pro-TGF-β1 and pro-activin A, with the exception of a five residue insert, which forms a short α-helix (lasso helix) in the latency lasso linking α1 and α2 helices. The α1 helix of the myostatin pro-domain occupies the major hydrophobic groove of one GF protomer, and is followed by the latency lasso which wraps between the ‘fingertips’ of the GF domain, providing an interface between pro and GF domains. The downstream α2 helix extends across the convex surface of one GF protomer, and occludes the type II receptor binding site (Figure 2A).

Electron density for the sequence linking the pro-domain forearm to the arm domain, and housing the TLD cleavage site (Arg98/Asp99), is missing in both protomers. Based on the distances between resolved residues and the directionality of electron density, it is apparent that the connectivity from the pro-domain forearm to the arm is such that the forearm interacts with the GF domain from the same chain, but with the arm from the opposite chain, giving rise to a domain-swapped arrangement, as is the case for pro-activin A (Figure 2B) ^12^. The distance from the last resolved residue of the forearm (Asp95) to the first visible residue of the arm (Glu107) is 22.7Å in our proposed domain-swapped arrangement (**Figure S5**). In the alternative connectivity, the missing 11 residues must span 35.2Å, which would require a near linear trajectory between endpoints. Such a constrained structural feature is unlikely given the lack of electron density in this region. With the domain-swapped topology, the extent to which the V-shaped dimer can open up will be limited by the linker sequence between α2-helix and the arm domain binding to the opposite mature domain. Our high resolution structure is missing 11 and 12 residues from this linker in the two protomers and the last visible residues are 23 and 24 Å apart, respectively, suggesting that a more open conformation could be still be accommodated (**Figure S5**).

Similarly, density for the furin cleavage site and sequence linking the pro-domain to the GF domain is weak and missing in places, however we were able to trace the entire main-chain connecting the pro-and mature domains in one of the two protomers. The density supports an additional domain-swapped conformation in which the pro-domain arm of one chain interacts with the GF domain of the other (**Figure S5**). It is noteworthy that the furin site is visible in our structure, as it suggests that pro-myostatin is a more constrained substrate for furin than pro-TGF-β1 and pro-activin A, which both lack density for the furin site. The uncleaved pro-activin A structure is missing ten residues with 16 Å between the last visible residues, whereas in pro-myostatin 12 residues span a direct distance of 34 Å resulting in a less flexible and therefore less accessible furin cleavage site (**Figure S5**).

Given the unusual open-armed conformation and lack of interaction between pro-domain arms, the question arises as to what drives the increased latency of the pro-myostatin complex over non-latent superfamily members.

### Latency conferring interactions of the pro-domain forearm

One of the key latency determining regions of the TGF-β superfamily pro-domains is the N-terminal helix-loop-helix forearm motif, with residues 42-115 originally identified as the inhibitory fragment of pro-myostatin^5^. This range incorporates the entire forearm region, extending from the N-terminus of the α1 helix to the start of the arm domain, and many of these latency conferring interactions are conserved between pro-myostatin and pro-TGF-β1^8^. In our structures, the α1 helix is clearly helical in nature from Arg45 to Leu64. As anticipated, the pro-domain α1 helix interaction with the GF domain is dominated by hydrophobic interactions, and of the seven aliphatic residues within this helical sequence, six are buried within the hydrophobic groove of the GF domain (Figure 4A). These aliphatic residues are conserved in pro-TGF-β1 (with exception of Ile58), and are known to contribute towards its latency (Figure 4F)^24^. Takayama *et al.* have synthesised a range of myostatin inhibitory peptides based on the mouse pro-domain α1 helix sequence, the best of which (Trp44-Leu64) binds to mature myostatin with a K_D_ of 29 nM and has been shown to increase muscle mass in mouse models muscular dystrophy^25^. The same authors have shown by alanine scanning that the aforementioned hydrophobic residues are critical to the inhibitory function of these peptides^26^. Nevertheless, the affinities of these α1 helix-derived peptides are not high enough to fully explain the latency, in line with the fact that many of these residues are conserved in non-latent pro-activin A (Figure 4F).

**Figure 4.**
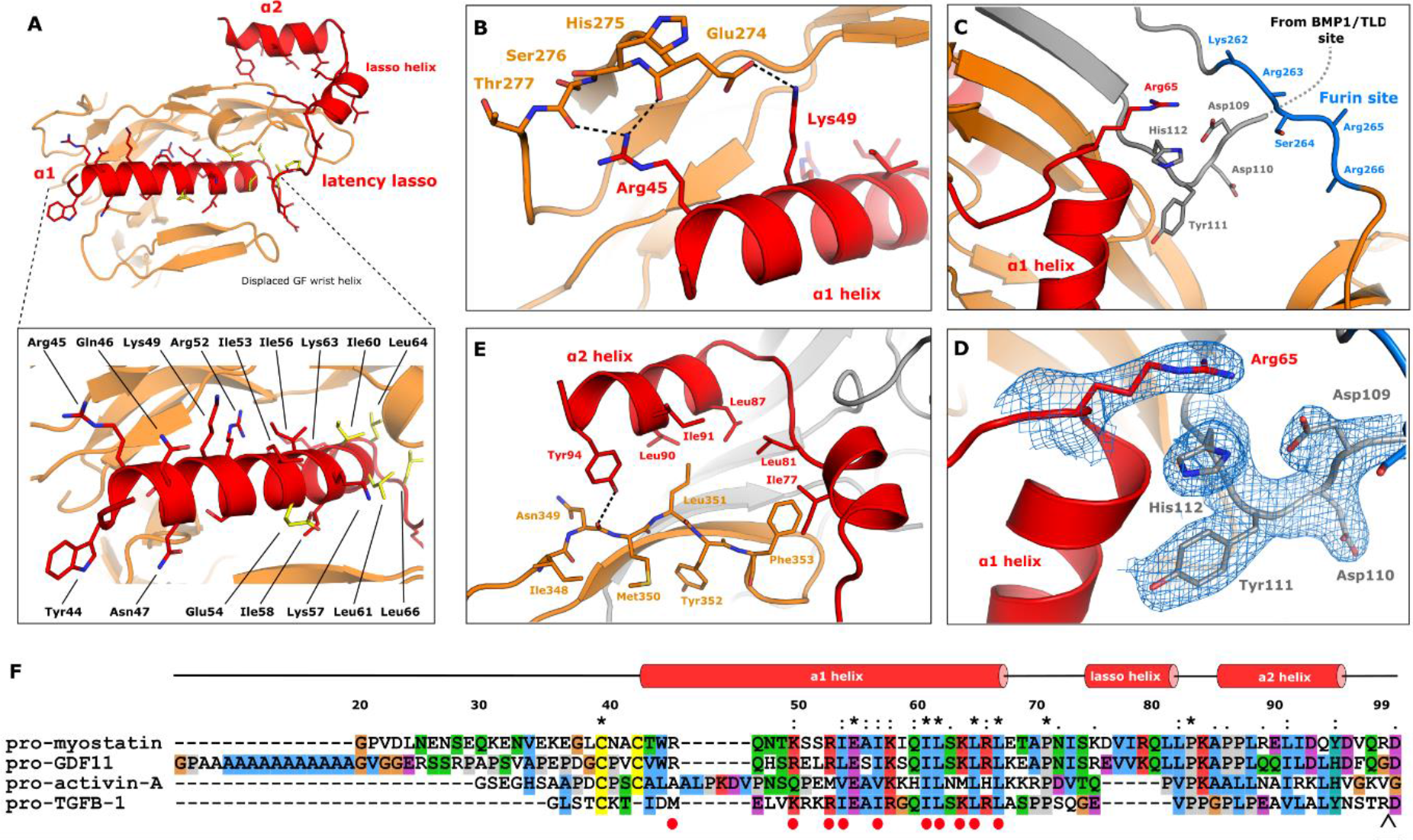
Key pro-domain interactions A. Interaction of N-terminal forearm of pro-myostatin (red) with mature GF domain (orange). Forearm residues within 4.5Å of mature GF are shown as sticks. Residues fully conserved between pro-myostatin, pro-TGF-β1 and pro-Activin A are coloured yellow. B. N-terminal α1 helix interactions with mature GF. C. Fastener residue interactions and proximity to the furin cleavage site (blue). D. Details of fastener stacking interaction, with electron density contoured to 1σ. E. α2 helix interactions with the convex surface of GF finger. F. Sequence alignment of N-terminal forearm regions (starting at first residue following signal peptide cleavage site). Alignment numbering and secondary structure annotation is based on the sequence and structure of pro-myostatin. Conserved residues are indicated by asterisks above alignment. Pro-myostatin α1 helix residues involved in GF interaction are indicated with red circles and the TLD cleavage site with a black arrowhead.

In addition to hydrophobic contributions, a number of electrostatic interactions appear to stabilise the α1 helix:GF interface. Arg45 forms hydrogen bonds with backbone carbonyls of Glu274 and Ser276 located on the N-terminal extension of the GF domain (Figure 4B), but truncation of the arginine sidechain has been reported to have little effect on the inhibitory function of α1 helix derived peptides^26^. Lys49 forms a salt bridge with the side chain of Glu274 while Arg52 forms multiple hydrogen bonds to backbone carbonyls of Ala306, Asn307 and Met367 from the GF fingers and Lys63, near the C-terminus of the α1 helix hydrogen H-bonds to the main-chain carbonyl of Pro365.

Cationic residues within the α1 helix of pro-TGF-β1 are reported to mediate non-covalent interaction with ECM bound LTBP-1, promoting subsequent covalent linkage through Cys33^24^. Given the conservation of these residues in pro-myostatin, and the prior observation that pro-myostatin interacts non-covalently with LTBP-3 (the primary LTBP expressed in skeletal muscle), it is possible that these interactions are conserved^13^. From a structural perspective, these charged residues may maintain the complex in a conformation that is competent for LTBP association (Arg52 and Lys63 are buried in the α1:GF interface), or form part of the LTBP-3 docking site (Lys57 is exposed to solvent and thus a potential LTBP-3 binding candidate).

The latency lasso extending from the C-terminus of the α1 helix wraps around the mature domain fingertips. A five amino-acid insertion in the latency lasso, unique to pro-myostatin and pro-GDF11, creates a short ‘lasso helix’ not observed in other pro-TGF-β family structures (Figure 4A). The downstream α2 helix lies against the convex face of the GF, occluding the putative type II receptor site. Tyr94 of the α2 helix forms a hydrogen bond to the Asn349 backbone carbonyl, an interaction also observed in pro-TGF-β1 (Figure 4E). The forearm:GF interface is dominated by aliphatic residues, and the shielding of these hydrophobic surfaces by the pro-domain is consistent with the vastly increased solubility of the pro-forms over the mature ligands, which are notoriously prone to aggregation under physiological-like buffer conditions.

### The pro-domain arm forms an extensive stabilising interface

The globular arm domain of pro-myostatin is structurally conserved with other pro-TGF-β superfamily structures and consists of two anti-parallel β-sheets and a short α-helix. Unlike pro-TGF-β1, the pro-myostatin arm domain lacks the β8/9 hairpin extension which facilitates covalent dimerization of the pro-TGF-β1 pro-domains^8^.

One of the distinguishing features of pro-myostatin is the substantial interface that the globular arm domain shares with the GF/forearm. The arm adopts a markedly different conformation to that of pro-activin A and pro-TGF-β1. Given the lack of a covalent constraint (as in pro-TGF-β1), the pro-myostatin arm is rotated almost ‘parallel’ to the mature domain, forms an extended β-sheet with the GF domain and creates a considerably larger interface with the GF and forearm helices (Figure 5A). This results in extended hydrogen bonding at the antiparallel β-sheet interface between GF β7’strand and arm β1 strand, with eight hydrogen bonds, compared to five in pro-Activin A, four in pro-BMP9, and only two for pro-TGF-β1 (Figure 5B). In the case of pro-TGF-β1, the reduced β-sheet interface is the result of considerable twisting between mature and arm domains to accommodate its ‘closed’ conformation.

**Figure 5.**
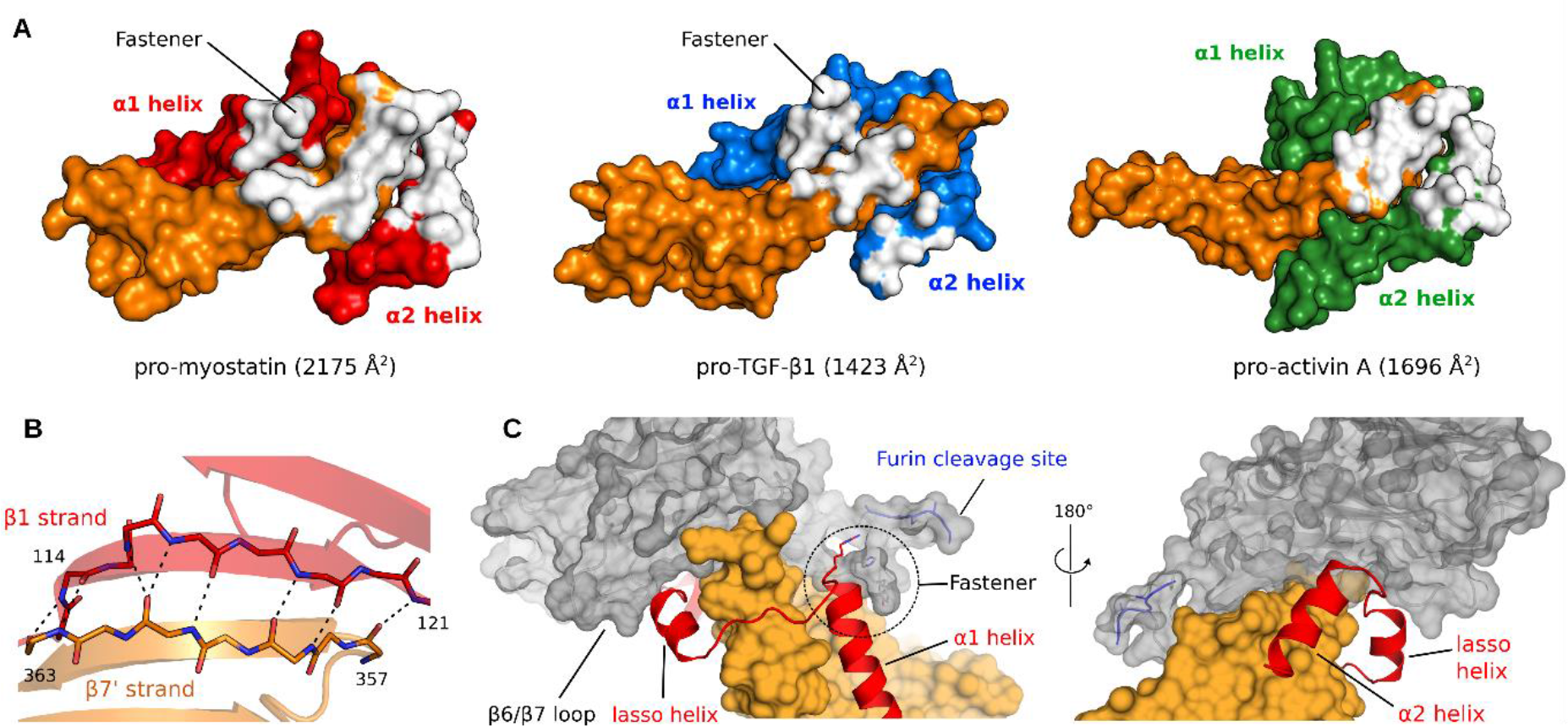
Pro-domain arm interactions A. Surface representation of mature GF protomer (orange) and pro-domain forearm (red, blue or green) showing the pro-domain arm interaction surface area (white, calculated using PyMOL). B. Extended hydrogen bonding (dashed lines) between the pro-domain arm (red) and mature domain (orange) with stick models of the two strands overlaid on cartoon representation of the same structure. C. The forearm (red) interaction with the mature GF (orange) is stabilised by interaction of the α1 helix with arm (grey) at the fastener, and by lasso interaction with loop β6/β7 of the arm domain. This binding mode completely encircles the GF finger, and occludes both putative receptor sites. For simplicity, only a single protomer is shown.

The pro-myostatin arm straddles the GF, and interacts with the forearm on both convex and concave sides of the GF, effectively sandwiching it between (Figure 5A). The β6/β7 loop of the arm domain latches over the latency lasso, completing the circle of pro-domain elements which enclose the second finger of the GF creating a straightjacket-like structure around it (Figure 5C). This extensive interaction may function to both mask surface hydrophobicity, and stabilise the furin cleaved complex, preventing spontaneous activation by dissociation. Furthermore, there is a well resolved stacking interaction between Arg65 at the tip of the α1 helix, with Tyr111 and His112 from the linker containing the TLD cleavage site, which appears to hold this cluster of charged features close to the α1 helix, further stabilising the arm/latency lasso interface (Figures 4C, 4D, 5C). These residues are equivalent to the ‘fastener’ residues described for pro-TGF-β1, non-conserved mutation of which was shown to liberate TGF-β1 signalling^8^.

It is interesting to note that the highly acidic sequence downstream of the TLD cleavage site (Glu107, Asp108, Asp109, Asp110) and the highly basic furin consensus sequence from the same chain almost overlap within the cavity between protomers (Figure 4C). It is tempting to speculate that interaction between the primary and secondary cleavage sites may play a role in the regulation of pro-myostatin activation. In the domain-swapped arrangement, in which the same chain crosses first from the N-terminal forearm to the arm domain on the opposite side and then back again to the mature GF, the entire complex is supported by criss-cross connectivity with the furin site coming over the TLD site, possibly providing steric protection of the latter. Cleaving the furin site would release the first of these tethers, potentially increasing lability of the pro-domain arms. For activin A, furin cleavage seems to be sufficient to release the latency that the pro-domain exerts, whereas in the case of myostatin, the arm domain interaction is strong enough to prevent dissociation from the mature growth factor, stabilised by the link from α2 helix to the first β-strand of the pro-domain. Subsequent cleavage of the TLD site removes this second tether, possibly disengaging the fastener interactions and allowing the arm, which is now attached only non-covalently, to dissociate from the GF/Forearm. The details of myostatin activation remain under investigation.

### Structural polymorphisms in human pro-myostatin

So far 134 unique naturally occurring missense mutations, involving 112 residues (77 in the pro-domain), have been identified in human pro-myostatin (Ensembl genome assembly GRCh38.p10, accessed on 05.06.2017). In order to further probe the molecular the determinants of latency, a series of pro-myostatin variants were made, designed either to recapitulate interesting natural polymorphisms, or to disrupt previously unappreciated interactions that we identified by structural analysis. In light of new structural information, we analysed the known polymorphisms and selected those that were most likely to affect pro-myostatin latency while eliminating residues that have already been shown to have an effect in previous studies (Table S2). We also selected several other residues that, in the absence of structural data, had not been analysed previously^25,26^. The most interesting substitutions for this analyses were those affecting the fastener (Arg65Ala, Arg65Cys, Tyr111His, His112Arg), and Lys153Arg, which has been associated with muscle and obesity related phenotypes^27–31^. In addition, we chose to analyse a naturally occuring Ala84Gly variant at the interface of forearm and arm domains as well as the role of Trp203 which forms part of the globular arm domain, and makes a hydrogen bond with backbone of Lys83 in the latency lasso using the indole nitrogen. Trp203 was mutated to Ala, His and Phe, in an attempt to minimise the effect of removal of the large side chain from the core of the arm. All mutated residues and their immediate surroundings are shown in Figure 6A.

**Figure 6.**
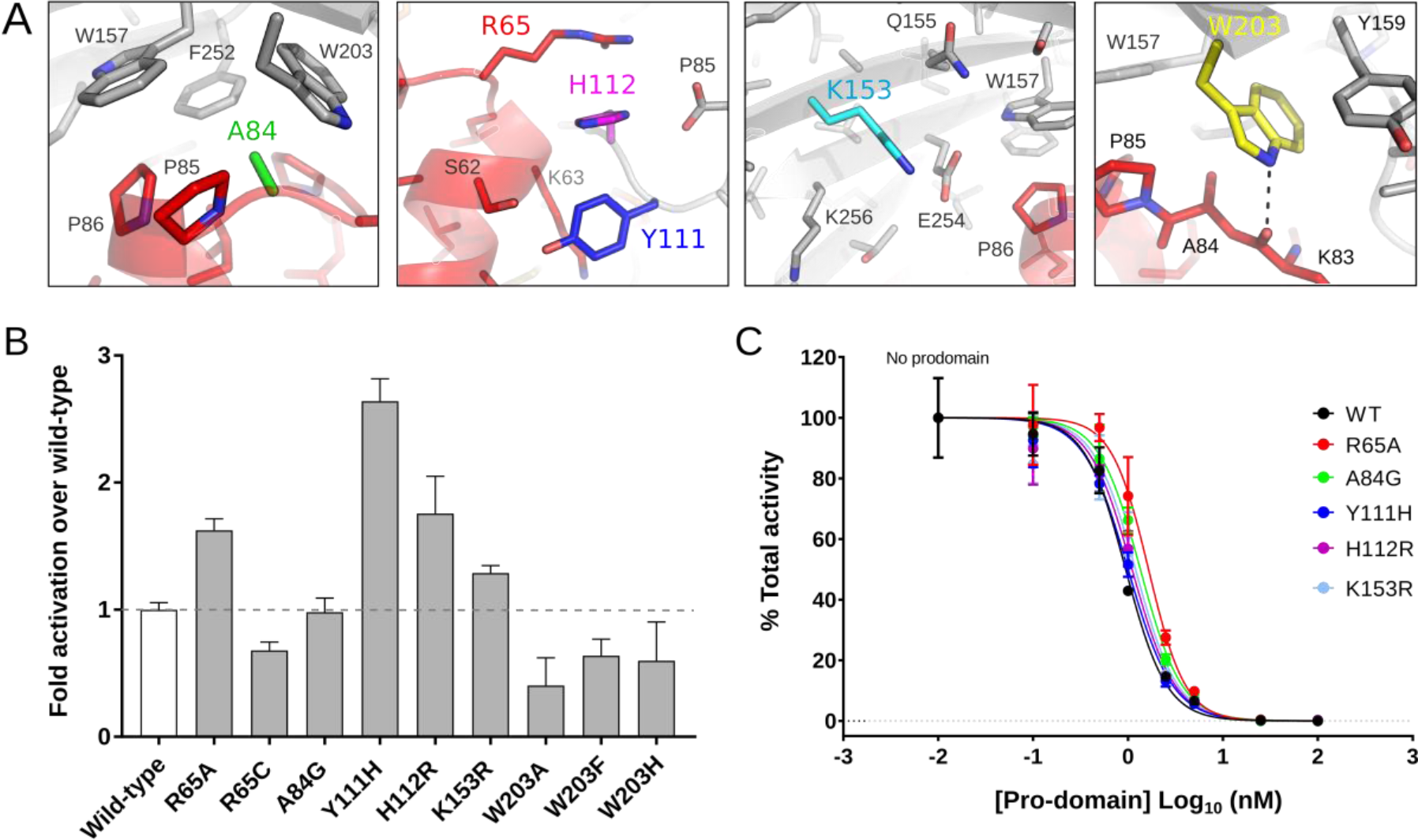
Pro-myostatin structural polymorphisms. A. Selected residues with known human missense polymorphisms and/or possible role in pro-mature interactions and latency of pro-myostatin. B. Signalling activity of pro-myostatin variants expressed in HEK293 cells with co-expression of furin and tolloid-like 2 proteases to facilitate activation (Data shown are means ± SEM). C. Inhibition of mature myostatin signalling by *E. coli* expressed pro-domains, determined using myostatin responsive luciferase reporter assay (in presence of 0.25 nM mature myostatin).

We first created expression constructs of selected mutants for production in HEK293-(CAGA)_12_ luciferase reporter cells, to analyse the effect of these mutations on bioactivity. In this setup, secreted pro-myostatin complexes showed minimal activity in the absence of cleavage at the tolloid-site, however when proteins were produced by co-transfecting cells with increased amounts of a construct encoding human tolloid-like 2, a number of variants showed deviation from wild-type activity levels (Figures 6B, S6). Mutation of Arg65 from the fastener motif to alanine increased signalling activity significantly over the wild type, but mutation of the same residue to cysteine abolished activity, presumably because of the detrimental effect of introducing a lone cysteine into an extracellular protein. The other fastener polymorphisms also increased myostatin activation with His112Arg behaving similarly to Arg65Ala, but Tyr111His was the most effective in reducing the latency of the protein, with over 2-fold higher activity compared to the wild type protein. Walker & McCoy *et al* similarly show that disruption of the fastener interaction (with Y111A, and H112A mutations) enhances activation of pro-myostatin over the wild-type^32^. The mechanism of increased activity of these variants is not certain, however it seems likely that weakening of the fastener interaction promotes increased dissociation of the pro-domain fragments following proteolysis by furin and TLD sites.

Somewhat surprisingly, the known polymorphic variant Lys153Arg had only a modest effect on myostatin activity, compared to wild-type protein^28^. The Ala84Gly variant showed no effect over the wild type protein. Mutations of Trp203 showed minimal increase in activity with low levels of tolloid co-expression, but at higher tolloid concentrations (where the wild-type protein was ca. 6x more active), Trp203 variants showed significantly reduced activity. This may be due to disruption of protein stability and/or folding and secretion, given that Trp203 is involved in a tightly packed hydrophobic interaction within the pro-domain arm.

To analyse the effect of these mutations in more detail, the same mutations were introduced into an *E. coli* expression construct of the myostatin pro-domain. The pro-domains (residues 24-262) were expressed solubly in *E. coli* as MBP fusions and assessed for their ability to inhibit mature myostatin in *trans*. Mutations of Trp203 and the Arg65Cys mutant gave very poorly soluble protein and thus were excluded from this part of the study. The wild-type myostatin pro-domain inhibited mature myostatin signalling in our experimental system, with an IC_50_ of 0.9 nM (95% C.I [0.83 – 1.08]) (Figure 6C). All variant forms of the pro-domain inhibited signalling with a similar range as the wild-type (Figure 6C, Table S4). The fact that these pro-domain variants did not recapitulate the same pattern of effects on activity observed for HEK293 expressed pro-complexes suggests these mutations do not meaningfully disrupt the pro:mature complex when reconstituted in *trans*. This may point to a mechanism in which the latency driving interactions are fully established only when the native protein folds and assembles into the domain-swapped complex. Alternatively, the effect of the mutations is in affecting the efficiency of proteolytic processing, without causing significant effect on the latency of the partially cleaved protein.

## DISCUSSION

Extracellular regulation of cell-signalling proteins is of clear biological importance, both during development and into maturity. Storage of signalling molecules in the extracellular matrix provides a means of rapid response to physiological change, avoiding the need to first synthesise, process and secrete the protein following stimulation. The mechanisms of extracellular storage and regulation of TGF-β family growth factors are diverse and a spectrum of latency exists within the pro-TGF-β superfamily, ranging from the fully auto-inhibited TGF-βs, to the BMPs and activins which readily dissociate from their pro-domains following cleavage, and instead rely on soluble antagonists to regulate signalling^33^. Pro-myostatin occupies an intermediate position on this scale, forming a weakly bioactive complex which requires further proteolytic activation to liberate its full signalling capacity^20^.

The pro-domains of TGF-β superfamily proteins are poorly conserved in sequence with comparison to the mature GF domains, making structural predictions and modelling of the pro-domains difficult^34^. In addition to variation at the amino-acid sequence level, the pro-forms for which we have structural information, show marked variability in their overall domain topology. The structures solved here show that latent pro-myostatin does not conform to the expectation of a ‘closed-arm’ conformation, as is seen for latent pro-TGF-β1. Instead, pro-myostatin forms an open, elongated structure, more reminiscent of non-latent pro-Activin A and pro-BMP9. The open-armed conformation observed crystallographically (in two distinct crystal forms) also exists in solution, as shown by SAXS, and interestingly, the overall conformation (and associated particle dimensions) does not seem to change significantly upon cleavage of the pro-domain. These findings are corroborated further by Le *et al*, who were able to clearly resolve the distinctive V-shape of pro-myostatin (and the furin cleaved complex), using negative stain electron microscopy^35^.

Despite differing overall topologies, the specific pro:GF interactions which drive the latency of these pro-complexes are mostly conserved with pro-TGF-β1, with the exception of the covalent linkage of TGF-β1 pro-domains at the bow-tie motif. In the absence of the avidity provided by the pro-domain dimerization, myostatin utilises other mechanisms to increase its latency compared to activin A, which in its overall topology is much more similar to myostatin. Our analyses suggest that there is no single critical feature that confers latency to the protein, but rather the latency arises from multiple features that increase the pro-mature affinity combinatorially.

We propose a model for the synthesis and activation of myostatin, based on the structures and data presented here and in other studies (Figure 7). With the domain-swapped, criss-crossed conformation of the protomer of pro-myostatin dimer, it is likely that a monomeric structure forms first, with the pro-domain supporting a dimerization-compatible conformation of the mature domain (Figure 7A-C). In this dimeric unprocessed precursor form of myostatin we can identify a number of features that contribute to its latency and provide a foundation for a tightly controlled activation process. The key features are: increased affinity of the α1 helix for the mature GF, a fastener epitope that locks the N-terminus around the mature GF domain fingers and the extended interface that the pro-domain arm makes with the mature GF and the latency lasso that binds to it, stabilising this complex (Figure 7D). This latency is released by a controlled, sequential proteolysis of the furin and TLD sites, with the furin site in particular being only moderately accessible and, at least before cleavage, partially obscuring the tolloid site (Figure 7E-F). Cleavage of the furin site alone is not sufficient for full activation, as the extended non-covalent interactions prevent the furin-cleaved complex from dissociating, even in the presence of competing high affinity antagonist follistatin. Release of the second TLD tether and separation of the two halves of the pro-domain is required before myostatin can exert its function. The arm domain is free to dissociate once the covalent linkage to the forearms is severed, which would then allow the helix-loop-helix epitope to dissociate as well (Figure 7G-H). Increased rates of hydrogen/deuterium exchange at pro:mature interaction sites following TLD cleavage as shown by Le *et al*, demonstrates increased lability of the shouder and forearm following TLD cleavage, priming the complex for dissociation^35^. Analysis of closely related GDF11 has shown that the N-terminal part of the pro-domain, with α1 and α2 helices can remain associated with the mature GF, promoting its solublity while not affecting bioactivity, consistent with the stepwise dissociation model^36^. Finally, removal of the α1 helix will enable the GF wrist-helix to form, re-establishing the type I receptor binding site (Figure 7I). The mature GF is now free to interact with its receptors and induce signalling. At the same time, this mature GF becomes a target for soluble inhibitors such as follistatin, which must act before the mature GF finds its receptor on cell surface.

**Figure 7.**
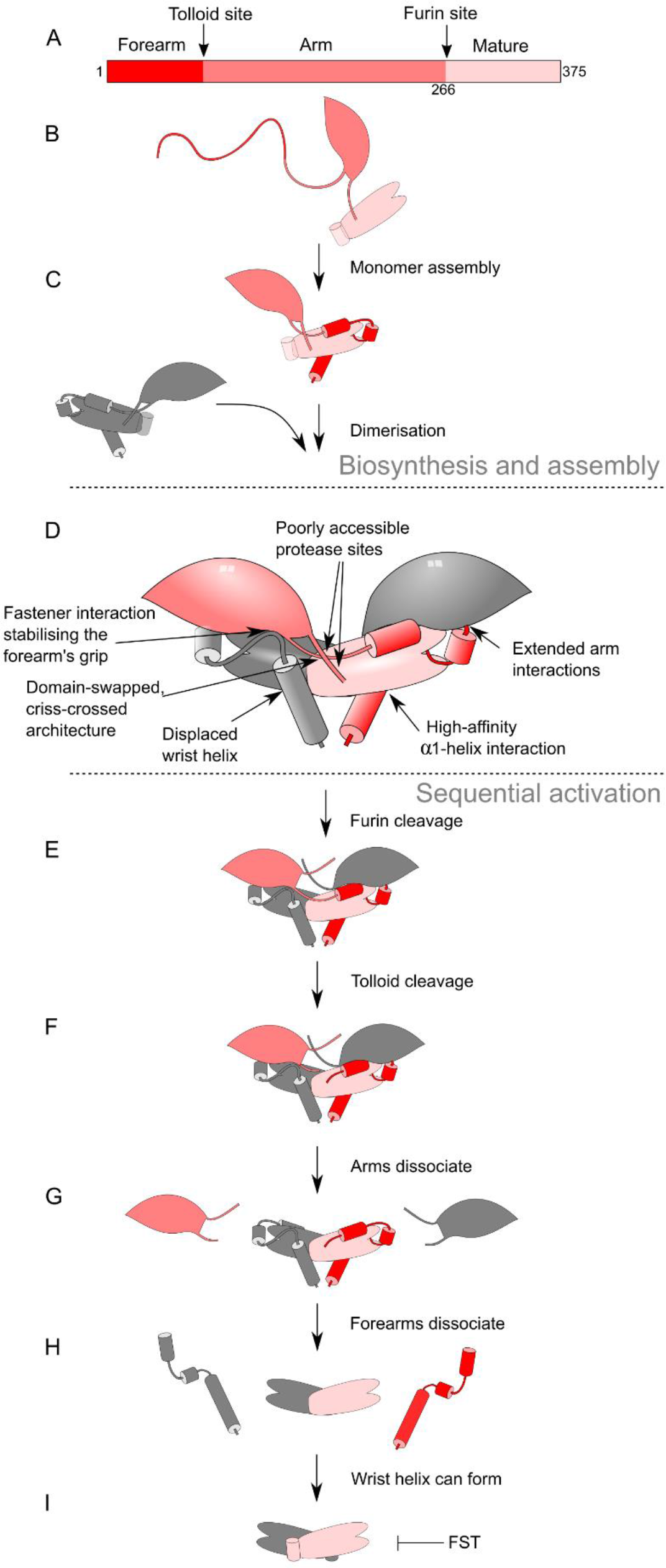
Model for myostatin biosynthesis and activation. The diagrams shows schematically different stages of myostatin biosynthesis (A-C), the features of the latent precursor (D) and the sequential activation of the pro-myostatin by furin and TLD (E-F), dissociation of the complex (G-H) and release of the mature GF (I).

It is possible that the fully latent complex can only assemble during synthesis, as supported by our mutagenesis data in which pro-domain variants which reduce latency of the pre-assembled complex do not have the same effect when the variant pro-domain is supplied in *trans*. This is consistent with the data of Walker & McCoy *et al*, who show that the pro-myostatin complex reconstituted from its individually purified components has reduced latency compared to the natively expressed complex.^32^ While our latent pro-myostatin complex shows low-level signalling activity in cellular assays, it is likely that the cleaved pro-complex is further stabilised *in vivo*, by interactions with components of the extracellular matrix, including perlecan and LTBP-3, which are known to bind elements of the pro-myostatin pro-domain^13,14^. Non-covalent bridging of pro-domains by ECM bound interactors could provide a mechanism for increasing functional affinity of the complex.

This structure and the analysis of the activation process provides us with a framework for assessing the effect of polymorphisms on myostatin function. Analysis of several missense polymorphisms in the pro-domain of myostatin demonstrate that some of these variants are more easily activated, and would potentially affect the musculature of those carrying the polymorphisms. As most of these polymorphism data are part of large scale anonymised studies, we are unable to correlate our results with phenotypic information relating to these individuals.

In addition to extending our understanding of the molecular details of myostatin activation, the crystal structures of this complex provide new ideas for the inhibition of myostatin signalling. A number of myostatin-neutralising antibodies have made it to clinical trials for a range of muscle wasting conditions, but these therapies tend to target the mature GF dimer which is present only for a limited period after activation, and shares significant structural homology with functionally unrelated TGF-β superfamily ligands. Stabilisation of the latent form can provide a more effective way for modulation of these growth factors and targeting the less conserved pro-region should increase the specificity of inhibition. These results will also facilitate future work in understanding other interactions pro-myostatin makes in the extracellular space and elucidation of the finer details of the activation process.

## METHODS

### Cloning and expression of pro-myostatin constructs for bacterial expression

A construct encoding human pro-myostatin lacking the signal peptide (residues 19-375, Uniprot 014793) was cloned into pHAT2 vector using *BamHI* and *NotI* restriction sites (pro-MSTN). The final construct contained an N-terminal His-tag and additional linker sequence, appending a total of 20 non-native amino acids to the protein N-terminus. For crystallography constructs, a TEV cleavage site was introduced into the N-terminal sequence by substituting the native sequence from Glu36-Cys42 with the TEV consensus ENLYFQGS, allowing removal of the predicted disordered N-terminus (proMSTN-Δ43) *in-vitro*^37^. Surface entropy reduction mutations, identified using the UCLA SERp server^21^ (G319A, K320A, K217A, Q218A, E220A), were introduced into the Δ43 crystallography construct (proMSTN-Δ43-mut).

For functional experiments, an additional construct was generated in which an HRV-3C protease site was engineered into the position of the native furin cleavage site, to allow robust cleavage *in-vitro* (proMSTN-3C). The aforementioned modifications to the original construct were completed using multi-step PCR protocol with overlapping oligonucleotide primers containing the modified sequence. The sequences of all oligonucleotides used for cloning are listed in Table S3.

Sequence-verified constructs were transformed into competent BL21(DE3)+pUBS520 cells by heat-shock and then grown overnight at 37°C overnight on LB-agar plates supplemented with 100 μg/ml of ampicillin and 25 μg/ml kanamycin. The cells were grown in 1L 2xYT media until OD_600_ between 0.8-1.0 and then for an additional 3 hours at 37°C after induction with 400 μM IPTG. The resulting cell pellet was harvested by centrifugation (4000 g, 20 mins).

### Refolding and purification of bacterially expressed pro-myostatin constructs

*E. coli* cells were lysed with Emulsiflex C5 and inclusion bodies prepared as per Wang *et al*, 2016^12^. The washed inclusion bodies, from 1L culture volume, were resuspended in 100 mM TCEP pH 7.0 and then solubilised by addition of 15 mL solubilisation buffer (8 M guanidine-HCl, 50 mM Tris-HCl pH 8.0, 10 mM EDTA and 0.1 M cystine) and incubated at room temperature while shaking, for 1 hour. The solubilised protein was clarified by centrifugation (15,000 g, 20 mins) and soluble material buffer exchanged into 6 M urea and 20 mM HCl, adjusted to 1 mg/mL and rapidly diluted 1:10 into 1L of cold refolding solution (100 mM Tris-HCl pH 9.0, 1 M pyridinium propyl sulfobetaine (PPS), 0.5 mM EDTA, 0.2 mM cystine and 2 mM cysteine) while stirring vigorously. The refolding solution was kept at 4°C for 7 days before purification.

One litre of refolding solution was filtered (0.65 µM Sartopure filter cartridge) prior to loading onto a 10 mL Source Q15 anion exchange column pre-equilibrated with 50 mM Tris-HCl pH 9.0. Five column volumes of the equilibration buffer were used to wash the unbound material, followed by elution with a linear gradient over 20 column volumes from 0-100% elution buffer (50 mM Tris-HCl pH 9.0, 1 M NaCl). For crystallographic constructs, the His-tagged N-terminus was removed by TEV protease cleavage. Following anion exchange capture, the pooled fractions were buffer exchanged to 20 mM Tris pH 8.0, 150 mM NaCl and incubated overnight with 200 µL TEV protease (2 mg/mL). TEV cleaved pro-myostatin was incubated with Ni-NTA resin (Cube Biotech) to separate the cleaved N-terminus and protease. After incubation with Ni-NTA resin for one hour at 4°C, the flow through containing cleaved protein was collected.

As a final step of purification for all constructs, protein fractions were concentrated and loaded onto HiLoad Superdex 200 16/60 gel filtration column (GE healthcare) pre-equilibrated with 20 mM Tris pH 8.0, 150 mM NaCl. Peak fractions were pooled and analysed by reduced and non-reduced SDS-PAGE.

Selenomethionine-labelled protein was expressed in minimal medium using metabolic suppression method to minimise endogenous methionine production and refolded and purified like the unlabelled protein. Selenomethionine incorporation was confirmed to be complete by mass spectrometry.

For production of the cleaved pro-myostatin complex, the native furin site was replaced by an HRV-3C cleavage site, and the protein purified as described for the wild-type protein, except for initial capture from refolding which was done using 5mL HiTrap Q HP (GE) column instead of Source Q15 column. The purified protein was then incubated with GST-HRV 3C fusion at a 4:1 mass ratio for 3 days at 4°C in gel filtration buffer (above). GST-tagged HRV 3C was separated from the cleaved complex by incubation with PureCube glutathione agarose resin (Cube Biotech). The complex was further purified by gel filtration (as above) with the dimeric mature domain and prodomain co-eluting as a single peak suggesting successful formation of a stable complex.

Mature myostatin was purified from the HRV-3C cleaved pro-myostatin complex by reverse phase chromatography (RPC). Acetonitrile and trifluoroacetic acid (TFA) were added to the purified complex, for final concentrations of 10% and 0.1% respectively. The acidified complex was then loaded onto ACE C8 300 4.6x250 mm RPC column, pre-equilibrated with 10% ACN and 0.1% TFA. The protein was eluted over 20 column volumes to 100% elution buffer (90% ACN, 0.1% TFA). Peak fractions were then dried by centrifugal evaporation. Mature myostatin was resuspended in 10 mM HCl prior to use. All protein concentrations were determined spectrophotometrically using calculated absorption coefficients at 280 nm.

### Bacterial expression and purification of wild-type and variant pro-domains

A cDNA fragment encoding the wild-type human pro-domain (residues 24-262, Uniprot 014793) was cloned into pET28a vector containing N-terminal 6x His tag and MBP fusion. To improve solubility and stability, the four pro-domain cysteines were mutated to serine and MBP was modified for surface entropy reduction according to Moon *et al*, 2010^38^. Specific polymorphisms were introduced into the pro-domain sequence using QuickChange PCR protocol with PfuUltra II Fusion HS DNA polymerase (600670, Agilent Technologies).

MBP-prodomain fusion constructs were transformed into competent Rosetta (DE3) Lac cells by heat-shock and then grown overnight at 37°C overnight on LB-agar plates supplemented with 34 μg/ml of chloramphenicol and 25 μg/ml kanamycin. The cells were grown in 1L 2xYT media until OD_600_ 0.6-0.8 and then overnight at 18°C after induction with 400 μM IPTG. The resulting cell pellet was harvested by centrifugation (4000 g, 20 mins) and resuspended in lysis buffer (20 mM Tris pH 8.0, 500 mM NaCl, 10 mM imidazole) supplemented with protease inhibitor (cOmplete Mini, EDTA-free, Roche). Following lysis with emusiflex C5, lysate was clarified by centrifugation (15,000 g, 20 mins), filtered and incubated with 1 mL Nickel-NTA resin (Cube Biotech) for 1 hour at 4°C. The resin was washed with 5x 5mL volumes of wash buffer (20 mM Tris pH 8.0, 500 mM NaCl, 20 mM imidazole) and eluted in 0.5 mL fractions with elution buffer (20 mM Tris pH 8.0, 500 mM NaCl, 500 mM imidazole). Nickel purified His-MBP-pro-domain fusions were concentrated and loaded onto HiLoad Superdex 200 16/60 gel filtration column (GE healthcare) pre-equilibrated with 20 mM Tris pH 8.0, 500 mM NaCl. Peak fractions were pooled and analysed by SDS-PAGE (Figure S7).

### Crystallisation and data collection

Purified unprocessed pro-myostatin crystallisation constructs were concentrated to 10 mg/mL and screened for crystallisation in commercial 96-well screens (Qiagen, Molecular Dimensions, Rigaku reagents). Sitting drops containing 300 nL protein solution and 150 nL reservoir were dispensed using Mosquito crystallisation robot (TTP Labtech), and incubated at 19°C. Subsequent optimisation screens were prepared in 96-well format using Dragonfly robotics (TTP Labtech), and sitting drops prepared as above.

Wild-type human pro-myostatin with N terminal truncation (proMSTN-Δ43), gave large (100-200 µm) cubic crystals in 0.1 M Na acetate (pH 4.2) with 1 M ammonium phosphate, reaching maximum size after one week. The subsequently engineered construct bearing the N-terminal truncation and combined surface mutations G319A, K320A, K217A, Q218A, E220A (proMSTNΔ43-mut) gave diffraction quality crystals overnight in 10% PEG 6K, 0.1 M HEPES pH 7.0. All crystals were flash frozen after transfer to a drop of mother liquor containing 26 % ethylene glycol.

### Structure determination

Diffraction data from cryo-cooled crystals were collected at Diamond Light Source on beamline I-03. For the SAD phasing of the SeMet-labelled proMSTN-Δ43, data were collected at the Se-peak wavelength (0.97970 Å) from multiple crystals, which had been grown under identical conditions. All data were processed using autoPROC (XDS, Pointless, Aimless, CCP4 suite) and the 7 highest quality datasets chosen for merging (Table S1)^39–43^. Their quality and mutual compatibility were assessed with regards to to diffraction quality, similarity of unit cell dimensions, resulting R_merge_ and quality of the anomalous signal. These datasets were merged using autoPROC/Aimless with a resolution cutoff of 4.19 Å; SAD phasing was performed using Phenix (AutoSol)^44,45^. The atomic model was built using Coot and the structure refined using phenix.refine and autoBUSTER^46^.

To determine the high-resolution structure of proMSTNΔ43-mut, data from a single crystal were processed using autoPROC to 2.59 Å. A partially refined low resolution model of proMSTN-Δ43 was used as a molecular replacement search model in PHASER, and model building and refinement were performed as above^47^.

Statistics of data collection, processing and refinement are shown in Table 1. Both the low and high resolution structures and their corresponding structure factors have been deposited in the Protein Data Bank (http://www.wwpdb.org/) with accession codes 5NXS and 5NTU, respectively. All structural figures were prepared using PyMol (Version 1.8 Schrödinger, LLC).

**Table 1.** Crystallographic data collection, processing and refinement statistics

|  |  |  |
| --- | --- | --- |
| Description: | Pro-MSTNΔ43-mut | Pro-MSTNΔ43 |
| PDB code: | 5NTU | 5NXS |
| **Data collection** |  |  |
| Synchrotron / beamline | DLS / I-03 | DLS / I-03 |
| X-ray wavelength [Å] | 0.97625 | 0.97970 |
| **Data processing \*** |  |  |
| Space group | C 1 2 1 | I 2 3 |
| Unit cell (a, b, c) [Å] | 168.16, 36.30, 120.45 | 196.83, 196.83, 196.83 |
| $\alpha, \beta, \gamma$ [°] | 90.0, 104.4, 90.0 | 90.0, 90.0, 90.0 |
| Resolution limits [Å]<sup>†</sup> | 76.26-2.58 (2.62-2.58) | 98.41-4.19 (4.27-4.19) |
| Number of protomers in ASU | 2 | 2 |
| No of total / unique reflections | 90386 / 22474 | 1517107 / 9470 |
| Multiplicity | 4.0 (4.2) | 160.2 (128.0) |
| R<sub>merge</sub> | 0.056 (0.826) | 0.141 (2.690) |
| R<sub>meas</sub> | 0.071 (1.037) | 0.148 (2.732) |
| I/σI | 11.7 (1.3) | 28.6 (3.0) |
| CC<sub>1/2</sub> | 0.997 (0.583) | 0.996 (0.923) |
| Completeness [%] | 97.7 (99.6) | 100.0 (100.0) |
| Anomalous completeness [%] |  | 100.0 (100.0) |
| Anomalous multiplicity |  | 84.0 (66.0) |
| Anomalous signal ( \|DANO\|/$\sigma$(DANO) ) | | 2.089 (7.368)<sup>‡</sup> |
| **Refinement** |  |  |
| R<sub>work</sub>/R<sub>free</sub> [%] | 0.215/ 0.260 | 0.274 / 0.301 |
| No. of unique/free reflections used | 22310/ 1118 | 9460 / 478 |
| R.m.s deviations: |  |  |
| bond lengths [Å] | 0.010 | 0.010 |
| bond angles [°] | 0.55 | 0.53 |
| Ramachandran analysis: (no. / % of residues) |  |  |
| Most favoured | 535 / 95 % | 452 / 88 % |
| Allowed | 23 / 4 % | 58 / 11 % |
| Outliers | 4 / 1 % | 5 / 1 % |
| Number of atoms/B-factors: |  |  |
| Protein atoms | 4404 / 100.5 | 3556 / 91.7 |
| Solvent atoms | 30 / 76.0 | 0 / - |
| Heterogen atoms | 86 / 90.8 | 0 / - |
| Mean/Wilson B-factor | 100.2/ 95.7 | 91.7/ 225.3 § |

\* Processing statistics are shown for data merged from 7 crystals
<sup>†</sup> Data in parenthesis are for the highest resolution shell.
<sup>‡</sup> Data in parenthesis are for the low resolution shell (98.41-11.38 Å)
§ The Wilson B-factor is ill-defined due to the low resolution of this structure

### SAXS data collection and analysis

Small angle X-ray scattering (SAXS) data were collected at the Soleil synchrotron SWING beamline (Gif-sur-Yvette, France) using a SEC-SAXS setup. The sample–detector distance was 1784 mm, providing a *q* range of 0.006–0.613Å^−1^ using a PCCD170170 (AVIEX) detector.

Samples (40 µL at a concentration of 7.5 mg/mL and 20 µL at 9 mg/mL for unprocessed pro-myostatin and cleaved pro-myostatin complex, respectively) were injected at 0.075 mL/min into a size-exclusion chromatography column (GE Superdex 200 Increase 10/300), pre-equilibrated with 20 mM Tris (pH 8.0) and 150 mM NaCl, in line with a quartz flow cell. Sample temperature was maintained at 293 K during data collection. 250 frames of scattering data were collected at an energy of 12000 eV during elution of each sample, with 0.75 s frame duration and 0.25 s dead time in between frames. In-house synchrotron software (FOXTROT 3.4.1) was used to select and average frames across elution peaks based on their R_g_ values and to subtract buffer scattering obtained from SEC flow-through data.

SCATTER 3.0 was used to plot scattered intensity (*I*) versus q for analysis of the forward scattering *I*(0) and radius of gyration (R_g_) from the Guinier approximation^48^. Guinier plots were linear for *qR*_g_< 1.3, suggesting samples were free of aggregation. DATGNOM (ATSAS package, EMBL) was used to calculate the pair-distance distribution function P(r), for estimation of maximum particle size (D_max_), based on truncated data-sets with *q-*ranges of 0.0063-0.2046 (unprocessed pro-myostatin) and 0.0114-0.2046 (cleaved pro-myostatin complex)^49^.

The *ab initio* modelling software DAMMIN (ATSAS, EMBL) was used to generate a molecular envelope of uncleaved pro-myostatin precursor. 34 independent *ab initio* models were generated, assuming P2 symmetry, averaged using DAMAVER (ATSAS, EMBL) and filtered by DAMFILT (ATSAS, EMBL) to give a final model. The crystal structure of unprocessed pro-myostatin (PDB code:5NTU) was docked into the envelope with SUPCOMB (ATSAS, EMBL) and visualised using PyMOL.

### Luciferase assay

In order to assess the signalling activity of purified pro and mature forms of myostatin, a dual-luciferase reporter assay using transiently transfected HEK293T cells (ATCC, catalogue no. CRL-3216; a generous gift from Dr Trevor Littlewood, Department of Biochemistry, University of Cambridge) was established. Cells were cultured (100 μl final volume per well) in 96-well flat-bottom cell culture plates using Dulbecco’s Modified Eagle Medium (DMEM; Life Technologies) with 10% (v/v) fetal bovine serum (FBS; Life Technologies) at 37 °C in a humidified incubator with 5% CO2. When the confluence of cells reached 80%, 33 ng of pGL3-CAGA (with myostatin responsive firefly luciferase reporter) and 17 ng of pRL-SV40 (Promega, with constitutively expressed *Renilla* luciferase) plasmids were mixed with 0.2 μl of FuGENE HD transfection reagent (Promega), and added to each well. 24 hours post-transfection, cell culture medium was removed, and replaced with DMEM containing 0.5% FBS and an appropriate dilution of myostatin, or one of its pro-forms. Each concentration point was repeated in triplicate. For pro-domain inhibition assays, purified wild-type and mutant variant MBP-prodomain fusions were serially diluted (0-100 nM) into DMEM containing 0.5% FBS and 0.25 nM mature myostatin, before adding to cells as above.

After overnight incubation in protein containing medium, cells were washed with PBS and lysed by addition of 20 μl Passive Lysis Buffer (Promega), and shaking at room temperature for 15 minutes. A volume of 4 μl of cell lysate from each well was transferred into a black flat-bottomed half-area 96-well plate. PHERAstar microplate reader (BMG LABTECH) was used to inject 15 µL of Firefly luciferase substrate (LAR II, Promega) per well, and measure resulting luminescence for 2 seconds after a 4-second delay. A volume of 15 μl of Stop & Glo Reagent (Promega) was then added into each well to quench the firefly luciferase signal and to provide substrate for the *Renilla* luciferase. *Renilla* luminescence measurements were measured as for Firefly luciferase. Firefly luminescence measurements were normalised against the *Renilla* luminescence. Non-linear curve fitting for EC_50_ and IC_50_ calculations were made using a variable slope (four parameters) dose response model in GraphPad Prism 7.

### Bioactivity assessment of pro-myostatin polymorphisms in HEK293 cells

HEK293 cells stably transfected with SMAD–responsive (CAGA_12_) luciferase-reporter gene were seeded at 20000 cells per well in 100 µl growth media into 96-well poly-D-Lys coated plates (655940 Greiner Bio-One GmbH, Germany) and grown until confluency of 75-85%. Cells were transfected with 25 ng pSF-CMV-FMDV IRES-Rluc bearing pro-myostatin variants, 50ng Furin DNA (pcDNA4) and 25ng human Tolloid-like 2 (pcDNA3 5) in OPTI-MEM reduced serum media (31985-070, Gibco, Life Technologies, USA). TransIT-LT1 Reagent was utilized for transfection (MIR 2300, Mirus Bio LLC, USA), 25µL transfection-reaction was added per well directly to the growth media and incubated (37°C, 5% CO2).

Six hours post transfection the media was removed and replaced with 100 µL serum-free media. 30 hours post transfection the cells were lysed using 20 µL per well 1x Passive Lysis Buffer (E1941, Promega, USA) with shaking (800 rpm, 20min, room temp.). Lysates were transferred to opaque black and white 96 well plates, and 40 µL of LAR II (Promega) was added, Firefly luminescence was recorded on Synergy H1 Hybrid Plate Reader (BioTek). Subsequently, 40 µL of Stop & Glo substrate (Promega) was added and *Renilla* luminescence was recorded. Firefly luminescence was normalized against *Renilla* luminescence. Signalling measurements for each pro-myostatin variant was repeated in triplicate, and the entire experiment run three times independently.

### Biolayer interferometry

To analyse the dissociation of mature growth factors from their pro-domains, biolayer interferometry (BLI) experiments were performed using ForteBio Octet RED96. As the pro-domains carry an N-terminal His-tag, the uncleaved pro-forms and cleaved complexes of pro-myostatin and pro-activin A were loaded onto the anti-penta-HIS (HIS1K) biosensors at the concentration of 20 µg/ml for 90 seconds. The immobilised biosensors were then immersed in kinetic buffer (PBS with 0.1% BSA and 0.02% Tween-20) with or without 500 nM follistatin-288 to observe the dissociation for 900 seconds. Follistatin-288 was expressed and purified as per Harrington *et al*, 2009. ^50^

### Size-exclusion chromatography multi-angle light scattering (SEC-MALS)

SEC-MALS analysis was conducted using Superdex 200 Increase 10/300 column (GE Healthcare) with DAWN HELEOS II light scattering detector (Wyatt Technology) and Optilab T-rEX refractive index detector (Wyatt Technology). Bovine serum albumin (Thermo Scientific) was used for calibration of the system in 20 mM Tris pH 8.0, 150 mM NaCl) before 100 μl of sample at a concentration of 1-1.5 mg ml^−1^ was analysed. Experimental data was recorded and processed using ASTRA (Wyatt Technology) software.

## Acknowledgements

We are grateful for the continuous access to the X-ray crystallography and Biophysics facilities at the Department of Biochemistry and for the support by the facility managers Drs Dimitri Chirgadze and Katherine Stott. We would like to thank Diamond (proposal mx14043, beamline i03), ESRF (proposal MX1727, beamlines ID30-A3 and BM14) and Soleil (proposal 20151251, SWING beamline) synchrotrons for the beamtime and support by the beamline managers, all of which contributed to the results obtained in this project. TC is funded by a Herchel Smith Scholarship.

## Author contributions

MH and TC conceived the study. TC produced the proteins, performed the biochemical, cellular and structural analyses and determined the crystal structures. GF solved the SeMet-labelled structure and refined the structures. GF and TC collected and processed the SAXS data. XW performed biophysical analyses and cell-based assays. JM, MG and TT designed and executed the analysis of myostatin mutants. MH and TC analysed the data and wrote the manuscript, with contributions from all authors.

